# Panacus: fast and exact pangenome growth and core size estimation

**DOI:** 10.1101/2024.06.11.598418

**Authors:** Luca Parmigiani, Erik Garrison, Jens Stoye, Tobias Marschall, Daniel Doerr

## Abstract

**Motivation:** Using a single linear reference genome poses a limitation to exploring the full genomic diversity of a species. The release of a draft human pangenome underscores the increasing relevance of pangenomics to overcome these limitations. Pangenomes are commonly represented as graphs, which can represent billions of base pairs of sequence. Presently, there is a lack of scalable software able to perform key tasks on pangenomes, such as quantifying universally shared sequence across genomes (the *core genome*) and measuring the extent of genomic variability as a function of sample size (*pangenome growth*).

**Results:** We introduce Panacus (pangenome-abacus), a tool designed to rapidly perform these tasks and visualize the results in interactive plots. Panacus can process GFA files, the accepted standard for pangenome graphs, and is able to analyze a human pangenome graph with 110 million nodes in less than one hour.

**Availability:** Panacus is implemented in Rust and is published as Open Source software under the MIT license. The source code and documentation are available at https://github.com/marschall-lab/panacus. Panacus can be installed via Bioconda at https://bioconda.github.io/recipes/panacus/README.html.

**Contact:** Luca Parmigiani, Daniel Doerr.

## 1 Introduction

The field of pangenomics emerged from the study of bacterial genomes (Tettelin *et al*., 2005), initially defining the pangenome as the complete set of genes within a species. This gene-based approach views the pangenome as the union of all genes across strains, distinguishing between the *core genome* – genes shared by all strains – and the *accessory genome* – genes found in one or more, but not all, strains.

Since the gene-based approach requires fully annotated genomes and excludes non-coding areas of the genome, an alternative definition of pangenome was proposed based purely on DNA sequences (Bentley, 2009). In contrast to the gene-based view, a sequence-based approach represents a pangenome as the set of all non-redundant genomic sequences, including coding and non-coding regions, and seamlessly extends to eukaryotic genomes with their complex and larger genomes.

Regardless, both perspectives treat the pangenome as a set, either of genes or DNA sequences, allowing for the examination of genomic variability and similarities. Two concepts have been central to address these aspects: *pangenome growth* and the *core curve* (Tettelin *et al*., 2005). Pangenome growth measures the expansion of total genomic content as additional genomes are sequenced. This process starts with a single genome and incrementally includes new genomes, expanding the collective genomic repertoire, whether defined by sequences or genes. Since the order of genome inclusion can affect the growth curve, pangenome growth is defined as the average over all possible genome orders. Similarly, the core curve illustrates the average size of the core genome as additional genomes are added.

As pangenomics progressed, so did the representation of pangenomes, by retaining the order of genomic sequences through a graph (Marcus *et al*., 2014). This evolution led to the adoption of *sequence graphs*, where nodes represent sequences shared across genomes, and edges indicate the consecutive presence of these sequences within a genome (Paten *et al*., 2017). Sequence graphs have been extensively used for the genome assembly problem (Compeau *et al*., 2011), where they allow the reconstruction of the original sequence from multiple reads.

On the other hand, sequence graphs are typically lossy, implying they can encode for a larger set of sequences than those they were initially constructed from, which makes them less attractive for a faithful representation of pangenomes. Augmenting a sequence graph with paths that correspond to the original sequences forms a *pangenome graph* (also referred to as variation graph (Garrison *et al*., 2018)), the primary target of our tool.

While plenty of tools are available to manipulate sequence graphs, for example gfatools (Li *et al*., 2020) or gfastats (Formenti *et al*., 2022), pangenome graphs are less represented. Moreover, with the increase in the amount of genomic data we need efficient tools to handle pangenome graphs.

Here, we introduce Panacus (pangenome-abacus), a tool designed for rapid extraction of information from pangenomes represented as pangenome graphs in the Graphical Fragment Assembly (GFA) format. Panacus not only efficiently generates pangenome growth and core curves but also provides estimates of the pangenome’s expansion with the addition of more genomes. In our study, Panacus is applied to generate the growth and core size for two pangenomes: HPRC-PGGB a draft human pangenome graph from the Human Pangenome Reference Consortium (Liao *et al*., 2023), with a focus on euchromatic and intragenic regions, and ECOLI-PGGB a pangenome of 50 *Escherichia coli* genomes (Garrison *et al*., 2023).

## 2 Panacus

Panacus is designed to count a variety of elements within pangenome graphs, including nodes, edges and base pairs that we will collectively refer to as *countables*. We define the *coverage* of a countable as the number of distinct paths that include that countable. For example, the coverage of the edge (`TC`, `A`) in Fig. 1(a) is four while the node `GG` has a coverage of one, despite the green path traversing the node twice. Users can choose any type of countable to generate and visualize its coverage distributions.

**Figure 1:**
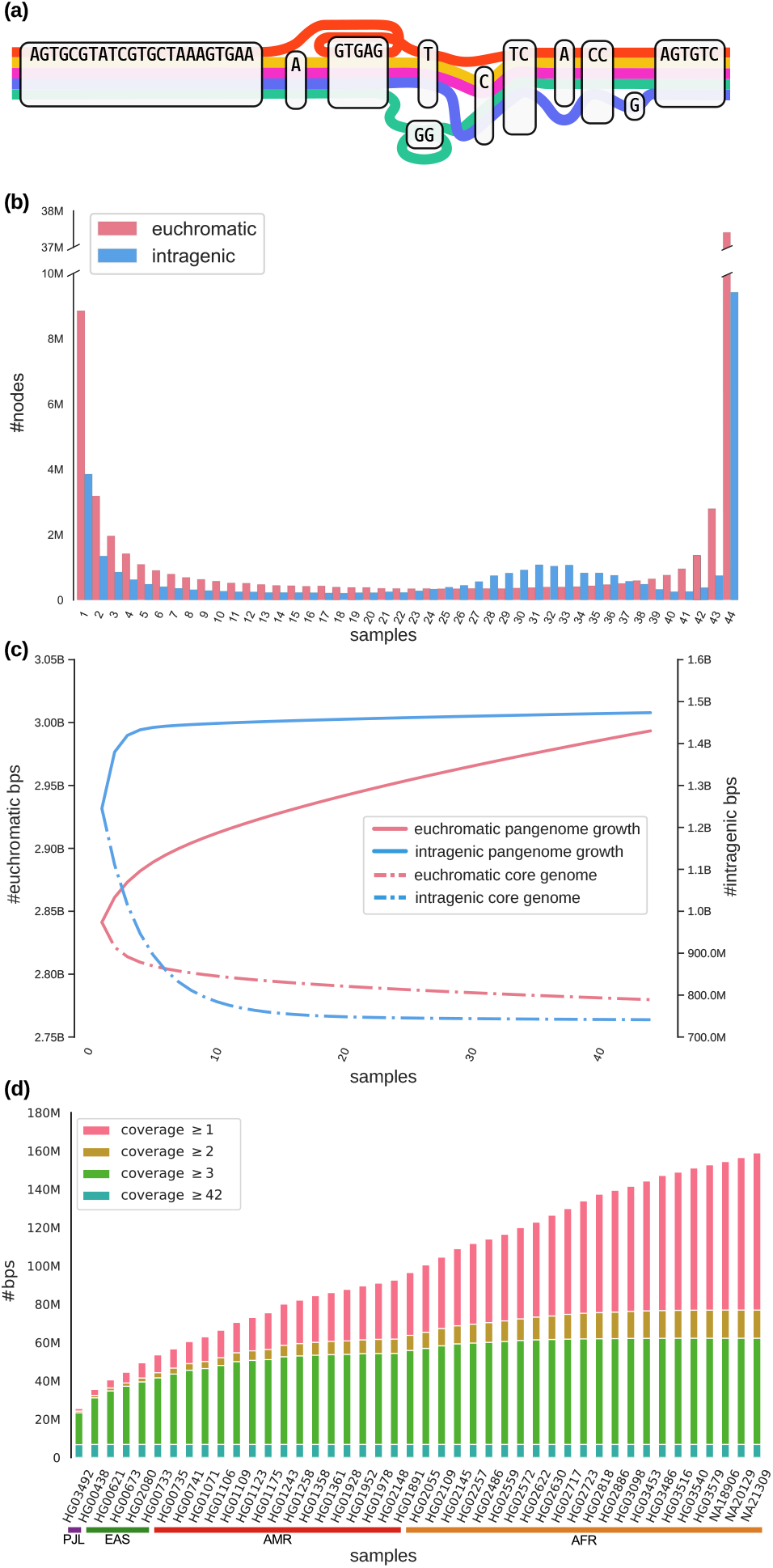
**(a)** A pangenome graph of five genomes.**(b)** Node coverage histograms of euchromatic and intragenic base pairs in HPRC-PGGB. **(c)** pangenome growth curves (solid) and core curves (dashed) of euchromatic and intragenic base pairs in HPRC-PGGB.**(d)** Ordered growth histogram re-created from (Liao *et al*., 2023, Fig. 3g) of the non-reference (GRCh38) euchromatic genome in HPRC-PGGB of 44 samples representing four populations, Punjabi from Lahore, Pakistan (PJL), East Asian (EAS), Admixed American (AMR), and African (AFR).

A key feature of Panacus is its ability to quickly calculate pangenome growth and core curves from the coverage frequencies of countables. Moreover, the tool allows to extract basic summary statistics or complete coverage tables for each countable. Finally, it produces an interactive report that contains the tabular data and its visualization within a standalone HTML page. Panacus is designed to take as input a file in GFA format^1^, where each line represents either a DNA segment (S-line), a link between two segments (L-line), or a path (P-line). Moreover, we support walks (W-lines), which were introduced in a successive version (v1.1) of the GFA format. In the subsequent text, we will not make any difference between P- and W-lines, but collectively refer to them as *paths*.

Since a path can represent multiple types of sequence, like a gene, a contig or even an entire chromosome, Panacus offers the option to group paths together. Users can group paths by passing a file containing the mapping or they can be automatically grouped by samples or haplotype following the PanSN notation^2^. All features that are subsequently presented support grouped paths.

Another feature of Panacus is the possibility to focus on, or exclude, specific parts of the pangenome. It allows selecting regions meeting a minimum coverage threshold or matching path names or path regions in a BED file. To mediate between the coordinate systems of paths and their corresponding DNA sequences, Panacus recognizes coordinate subsetting also in path names of P-lines, e.g., `path-id:10-200`.

A possible approach to construct pangenome growth curves involves creating sets of countables for each path, sampling a random order of the paths, and sequentially including them. This process is repeated for several permutations of the order of paths, taking the average of the total number of new countables at each inclusion step and obtaining an approximate expected growth of the pangenome.

In contrast, Panacus adopts a method similar to the one presented in Parmigiani *et al*. (2024) to efficiently calculate the pangenome growth and the core curve. This approach is both *exact* and efficient. It avoids sampling and repetitive computation of the union of paths, thereby saving both memory and time. Notably, this concept is not new and has been recognized in ecology since the 1970s, but it remains largely underutilized in bioinformatics. Parmigiani *et al*. (2024) applied this concept to show how *k*-mers (sequences of length *k*) found in multiple genomes yield a pangenome growth similar to that of genes. Here, Panacus uses this method to efficiently generate pangenome growth and core curves for countables within the paths of pangenome graphs.

The principle behind this approach is as follows: pangenome growth can be equivalently defined as either the average size of the incremental union of paths across all possible orders or the average size of the union of all combinations of *m* paths, where *m* goes from 1 to the total number of paths. According to the second definition, a countable contributes to the average each time the union of paths includes *at least* one of the paths that contains the countable. This means that the greater the coverage of a countable – the more paths are passing through it – the more it contributes to the average, since it is present in more combinations of *m* paths. The contribution of a countable can be calculated by subtracting from the number of combinations of *m* paths how many *do not* contain the countable. For example, if *m* = 3 and there are 10 paths with only 4 containing the countable, there are 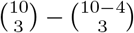 combinations of paths where at least one 3 3 of these four paths is present.

The same strategy applies to calculating the core curve, counting the contribution of a countable only if it is present in all *m* paths. Additionally, the definition of the core genome can be relaxed to encompass countables that are present in a certain percentage of paths. This percentage is determined by the user-defined *quorum* parameter, *q*, which can range from 0 to 1, specifying the proportion of paths in which a countable must be present to be considered part of the core.

A key data structure of the algorithm is the histogram *h*(*i*), as depicted in Fig. 1(b), which tracks the *coverage frequency* of the pangenome graph. This histogram stores the number of countables that appear in exactly *i* paths and it can be used to directly calculate the expected growth and core curves for each subset size *m*.

While accurately extrapolating pangenome growth and core genome size for new, unseen genomes is challenging, Panacus aids in making an educated estimate. To facilitate this, we provide a Python script within Panacus that enables users to extrapolate the growth and core curves using both parametric methods, such as power law (Tettelin *et al*., 2008) or exponential decay (Tettelin *et al*., 2005) and a non-parametric method, the Chao2 estimator (Chao, 1987).

## 3 Results

We study the pangenome growth of the draft human pangenome graph `hprc-v1.0.pggb` (HPRC-PGGB) recently released by the *Human Pangenome Reference Consortium* (Liao *et al*., 2023), and graph `ecoli50` (ECOLI-PGGB) that was published in Garrison *et al*. (2023). Both graphs have been constructed with *PanGenome Graph Builder* (pggb) (Garrison *et al*., 2023). The former graph constitutes the complete haplotype-resolved assemblies of 44 samples and reference assemblies *GRCh38* and *T2T-CHM13v1*.*1*, and encompasses overall 8.41 gigabases distributed over 111 million nodes. The latter graph is composed of genome assemblies from 50 *Escherichia coli* strains and consists of over 18 megabases and 1.5 million nodes.

We benchmark Panacus against a competing tool called `odgi heaps` (Guarracino *et al*., 2022) that is able to process pangenome graphs stored in GFA v1.0 format. This tool’s method is based on sampling permutations and as such produces only an estimate of the growth curve. Obtaining stable estimates of pangenome growth with `odgi heaps` depends on the number of generated samples. This affects the running time, which is further prolonged by the preceding construction of an index of the GFA file, unless already present. For these reasons, it is difficult to devise a fair comparison between the two methods. Table 2 reports the running time and memory consumption for both methods on the two graphs. We ran `odgi heaps` twice, once sampling only one permutation and another time sampling 100 permutations. While a single sample is generally insufficient to give a stable estimate, an average over 100 samples results in an acceptable estimation of the pangenome growth curve for the dataset at hand.

For HPRC-PGGB, the computation time for the pangenome growth step with `odgi heaps` is approximately 24 minutes, notably less than half the time required by Panacus, which takes nearly 55 minutes. However, when including the time for `odgi`’s graph indexing (around 5 hours), the total computation time for `odgi heaps` increases significantly, making it about 5 times slower than Panacus. Additionally, when `odgi heaps` processes 100 permutations, its completion time extends dramatically to nearly 47 hours, becoming a major time-consuming step. In terms of memory usage, `odgi` requires between 148 and 182 GB, whereas Panacus is much more memory-efficient, needing only 51 GB, which is approximately a third of `odgi`’s requirement. These trends are consistent with the benchmarks on the ECOLI-PGGB graph, where the over-all running times for `odgi` with 100 permutations, indexing included, accumulate to 11:55 minutes and for Panacus to 13 seconds.

Panacus also provides additional functionality not offered by `odgi heaps`, such as quantifying countables other than base pairs, and the simultaneous calculation of multiple other statistics, including core curves. A comparison of the features between `odgi heaps` and Panacus is shown in Table 1.

**Table 1:** Comparison between methods.

|  | Parmigiani *et al.* (2024) | odgi heaps | Panacus |
| --- | --- | --- | --- |
| Input format | FASTA | GFA v1.0/OG | GFA v1.1 |
| Input type | sequences | pangenome graph | pangenome graph |
| Countables | $k$-mers | bps | nodes/edges/bps |
| Growth calculation | exact | sampling | exact |
| Core curve calc. | exact | ✗ | exact |
| Quorum core curve calc. | exact | ✗ | exact |
| Ordered growth calc. | ✗ | ✗ | ✓ |
| Path grouping | ✗ | ✓ | ✓ |
| Coverage thresholding | ✗ | ✓ | ✓ |
| Subsetting | ✗ | ✓ | ✓ |
| Exclusion | ✗ | ✗ | ✓ |

**Table 2:** Comparison of CPU time (h:mm:ss) and Maximum Resident Set Size (GB) between Panacus and odgi heaps in computing pangenome growth curves on base pairs for HPRC-PGGB and ECOLI-PGGB.S.

| Dataset | Tool | CPU Time | Memory | Method |
| --- | --- | --- | --- | --- |
| HPRC-PGGB | **odgi build** | 5:01:56 | 182.41 | indexing |
|  | **odgi heaps** | 0:23:46 | 151.72 | 1 permutation |
|  |  | 47:13:54 | 148.89 | 100 permutations |
|  | Panacus | 0:54:40 | 51.03 | exact calculation |
| ECOLI-PGGB | **odgi build** | 0:49 | 1.53 | indexing |
|  | **odgi heaps** | 0:07 | 0.64 | 1 permutation |
|  |  | 11:06 | 0.64 | 100 permutations |
|  | Panacus | 0:13 | 0.36 | exact calculation |

Panacus’ capabilities of subsetting pangenome graphs enables direct analysis of specific region sets, rendering tedious manipulation of the graph’s GFA file unnecessary. We demonstrate this functionality by comparing the pangenome growth of euchromatic and intragenic regions of the human pangenome. By restricting to euchromatic sequence, we seek to avoid any error in our analysis caused by under-alignment. Intragenic regions, which are generally located within the euchromatin, are overall strongly conserved and contrast the much larger euchromatic sequence. Here we use coordinates of euchromatic and intragenic (incl. introns) regions provided by Liao *et al*. (2023) to calculate node coverage histograms (Fig.1(b)), and base pair-based pangenome growth and genome core curves (Fig. 1(c)). The intragenic sequence of HPRC-PGGB displays a non-convex coverage pattern caused by an amplified bias resulting from a lower proportion of shared sequence on the sex chromosomes, and quickly converging pangenome growth and core curves compared to those of the euchromatic sequence. The latter, despite having a higher ratio of shared nodes compared to unique nodes (Fig. 1(b)) and base pairs (not shown), displays a much steeper growth curve indicating a relatively higher degree of variability in the genomic sequences of the human pangenome at hand. At last, Panacus can also produce ordered growth histograms and was used to illustrate the growth of non-reference, euchromatic autosomal sequences of the human pangenome in (Liao *et al*., 2023, Fig. 3g), a recreation of which is shown in Fig. 1(d).

## 4 Conclusion

We have introduced Panacus, a versatile and efficient tool designed for exploring and comparing pangenome graphs. Panacus enables the rapid generation of pangenome growth and core curves directly from GFA files. In this paper, we demonstrated its use in comparing distinct areas within a human pangenome graph. Beyond this application, Panacus can also be employed to compare the same pangenome constructed using different tools, or to analyze completely different pangenomes, assessing their growth and core sizes.

## Acknowledgements

We thank Heng Li for helpful discussions.

## Footnotes

1 https://github.com/GFA-spec/GFA-spec

2 https://github.com/pangenome/PanSN-spec

## References

Bentley, S. (2009). Sequencing the species pan-genome. Nature Reviews Microbiology, 7(4), 258–259.

Chao, A. (1987). Estimating the population size for capture-recapture data with unequal catchability. Biometrics, 43(4), 783.

Compeau, P. E. C. et al. (2011). How to apply de Bruijn graphs to genome assembly. Nature Biotechnology, 29(11), 987–991.

Formenti, G. et al. (2022). Gfastats: conversion, evaluation and manipulation of genome sequences using assembly graphs. Bioinformatics, 38(17), 4214–4216.

Garrison, E. et al. (2018). Variation graph toolkit improves read mapping by representing genetic variation in the reference. Nature Biotechnology, 36(9), 875–879.

Garrison, E. et al. (2023). Building pangenome graphs. bioRxiv.

Guarracino, A. et al. (2022). ODGI: understanding pangenome graphs. Bioinformatics, 38(13), 3319–3326.

Li, H. et al. (2020). The design and construction of reference pangenome graphs with minigraph. Genome Biology, 21(1), 265.

Liao, W.-W. et al. (2023). A draft human pangenome reference. Nature, 617(7960), 312–324.

Marcus, S. et al. (2014). SplitMEM: a graphical algorithm for pan-genome analysis with suffix skips. Bioinformatics, 30(24), 3476–3483.

Parmigiani, L. et al. (2024). Revisiting pangenome openness with k-mers. PCI Community Journal, 4, e47.

Paten, B. et al. (2017). Genome graphs and the evolution of genome inference. Genome Research, 27(5), 665–676.

Tettelin, H. et al. (2005). Genome analysis of multiple pathogenic isolates of Streptococcus agalactiae: Implications for the microbial “pan-genome”. Proceedings of the National Academy of Sciences of the United States of America, 102(39), 13950–13955.

Tettelin, H. et al. (2008). Comparative genomics: the bacterial pan-genome. Current Opinion in Microbiology, 11(5), 472–477.

